# The traits of skull anatomy in green frogs of the genus *Pelophylax* (Anura, Ranidae)

**DOI:** 10.1101/2022.03.22.485387

**Authors:** Slutska Natalia

## Abstract

During dissecting skulls of green frogs of the genus *Pelophylax* it has found that the description of a number bones and cartilaginous structures requires addition and clarification. In addition, previously undescribed details of anatomical structure of the skull of green frogs has revealed. The traits of anatomical structure of skull of green frogs, characteristic for ancestral forms of Anura found too.

## Introduction

European aquatic green frogs - *Pelophylax. ridibundus* (Pallas, 1771) and *Pelophylax lessonae* (Camerano, 1882) are classic experimental objects for comparative anatomical, physiological and embryological researches (Gurtovoi N.N. et al., 1978; Dzerzhinsky F.J. et al., 2013). The frogs of the genus *Pelophylax* Fitzinger, 1843 are widely used in ecological studies because with other groups of tailless amphibians they are good indicators that appears sensitive to the effects of various environmental factors (Peskov V.N., Kotserzhinskaya I.M., 2004).

Tailless amphibians exhibit a vast range of cranial morphologies, life history strategies and ecologies (Bardua C. et al., 2020). Green frogs of the genus *Pelophylax* are typical representatives of tailless amphibians and play an important role in study of the evolution of vertebrates. With a number of traits of modern land vertebrates, European green frogs possess features of skull structure characteristic for paleozoic amphibians and ancient transitional forms from lobe-finned fishes to tetrapods. One of the basic body systems, that needed changes for life on land, is the skeleton, in particularly the skeleton of limbs and skull (Markov A., Naimark E., 2014; Shubin N., 2017). To understand the mechanisms of morphological evolution of the skull during the transition to terrestrial pattern of life, it is necessary to know well the skull anatomy of recent species of tailless amphibians (Konstantinov V.M., Shatalova S.P., 2005). Besides knowledge of anatomical structure and evolution of the skull of tailless amphibians helps to better understanding of basic evolutionary mechanisms underlying the development of the skull of representatives of higher classes terrestrial vertebrates (Swensson M., Haas A., 2005). It is also necessary for correct explanation of the results of experiments on the frogs and for identification of paleontological material (Böhme G. & Günther R., 1979).

During of the research we established that some very important traits of anatomical structure of the skull of frogs of the genus *Pelophylax* studied not enough. There are significant discrepancies in their description by different authors or the description of certain bone and cartilaginous structures requires clarification. Also revealed new details of skull structure not previously described in the literature, for example in the structure of upper jaw, lower jaw and in jaw joint or jaw articulation.

## Material and methods

The study conducted on the skulls of 100 adult *P. ridibundus* and on the skulls of 50 adult *P. lessonae*, caught in period 2003-2017 years in the Volhynia, Kiev, Odessa, Kharkov and Kherson regions.

To obtain the skulls, the heads of adult frogs fixed in 10% formalin, washed in water and removed from them eyes and skin. Then the heads put in 15-20% solution of sodium hypochlorite for 7-8 minutes to resolved soft tissues, but so that cartilaginous structures of the skull were preserved. Thereafter the remnants of soft tissues cleaned using surgical instruments (scissors, tweezers, dissecting needles). Then the skulls for studying of cartilaginous structures were stored in tap water at refrigerator. To research bone structures and permanent storage, the skulls degreased for a one day in white spirit, then they bleached for 2–3 days in a 3% hydrogen peroxide solution and dried on air at room temperature.

In this paper anatomical terminology applied from the works of Terentyev P.V. (1950), Špinar Z.V. (1972), Roček Z. (2003). The MBS-9 laboratory binocular magnifier used in this study.

## Results and discussion

The *Pelophylax* skull is square or rectangular, rather flattened and no have obvious ornamentation. The length of the orbit is nearly half of the total cranial length. The skulls of aquatic green frogs have maximum width at the posterior ends of lower jaw. Each nasal bone *(nasale)* coats about half of the olfactory capsule and usually narrowly separated at midline. Teeth are present on the *premaxilla, maxilla* and *vomer*. The squamosals are well developed, with a long zygomatic ramus that extends almost until pterygoid process of *pterygoideum*. The quadratojugals are slender. At the posterior ends of jaw’s joints there are visible areas of cartilage (cartilaginous part of *pars quadrata palatoquadrati*). New traits of skull anatomy *Pelophylax* frogs found by us and described in this article presented in the scheme below (Fig. 1).

**Fig. 1.**
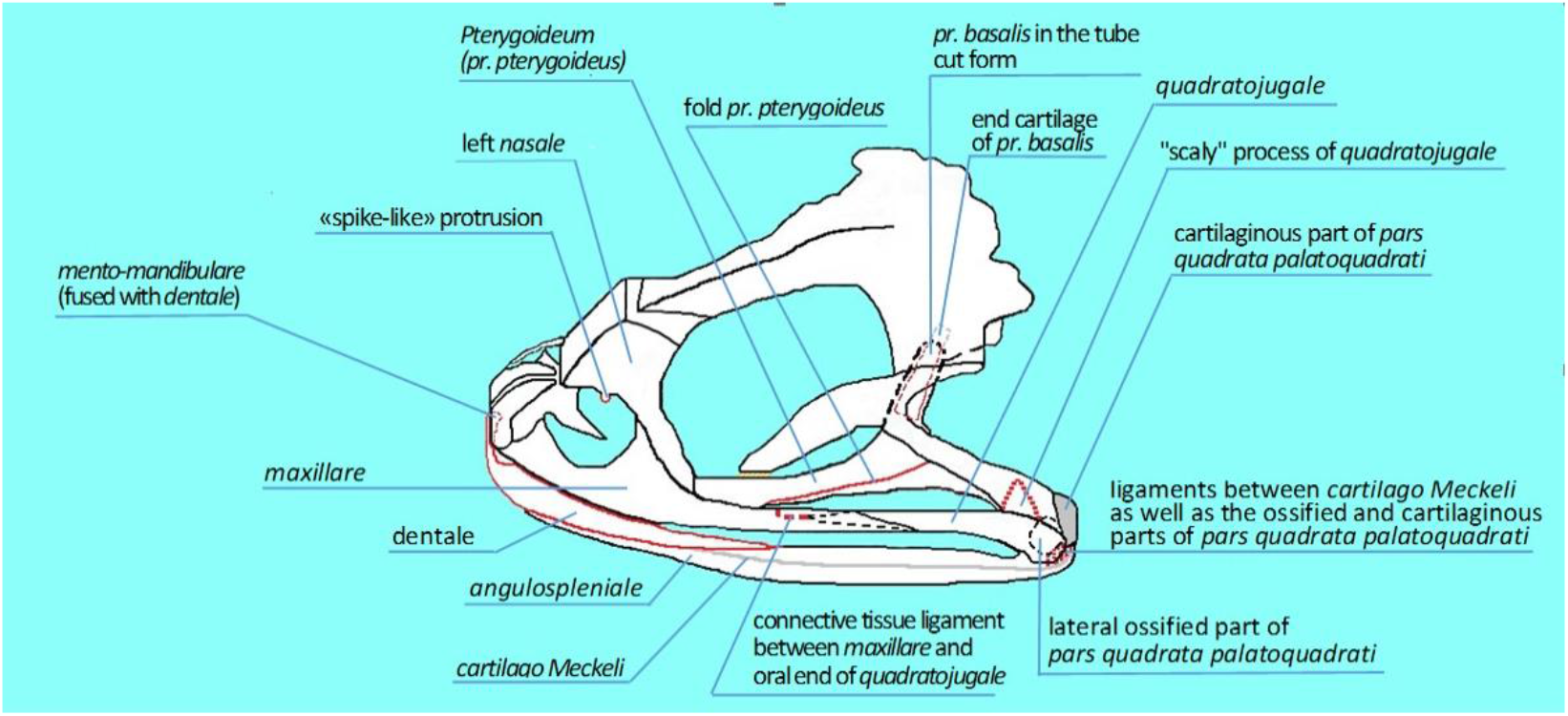
Scheme of adult *Pelophylax* frog skull, lateral view. New traits of anatomy, described in this article. No scale.

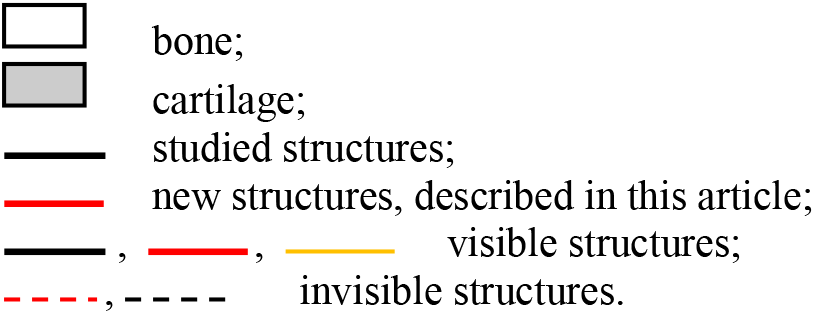

### Nasal bones *(os nasale)*

*Nasale* are part of skull roof of tailless amphibians and appear integumentary bones. Their shape was more than once varying in phylogenesis of Anura. For example, in Jurassic frogs *Notobatrachus degiustoi* Reig, (Baez A., Nicoli L. 2004) *nasale* were wide, had an irregular shape, and did not have lateral branches (*ramus lateralis*) (Špinar Z.V., 1972). In contrast, in frogs *Eopelobates bayeri* Špinar, 1952 (Anura, Pelobatidae) from Central Europe, found in the oligomyocene layers in Bohemia (Czechoslovakia), *nasale* had an exact triangular shape with pronounced *ramus lateralis*. In extant tree frogs from Argentina *Hypsiboas pulchellus* (Anura, Hylidae) *nasale* have a crescent shape and through *ramus lateralis* come to contact with facial parts *(pars facialis)* of the maxillary bones (Hoyos J.M. et al., 2012). In *Gastrotheca gemma* sp. nov. (Anura, Gastrotheca), new species was discovered in northeastern Peru, *nasale* have quadrangle shape (Venegas P.J. et al., 2021).

During dissecting frog skulls *P. ridibundus* and *P. lessonae* we found that *nasale* do not have a triangle shape as described by many authors (Terentyev P.V., 1950; Gurtovoi N.N. et al., 1978; Kartashev N.N. et al., 2004). At the base of their *ramus lateralis* there are clearly visible sharp bony protrusions, which we called “spike-like” protrusions (Fig. 2), which are not mentioned by other authors when described *nasale* of green frogs. As well as the “spike-like” protrusions of *nasale* not described in ancient and extant representatives of Anura mentioned above. We can assumed that form of *nasale* with “spike-like” protrusions in studied extant representatives of genus *Pelophylax* appears one from variants of structure of these bones in species of Anura and may be result of further reduction of *nasale*.

**Fig. 2.**
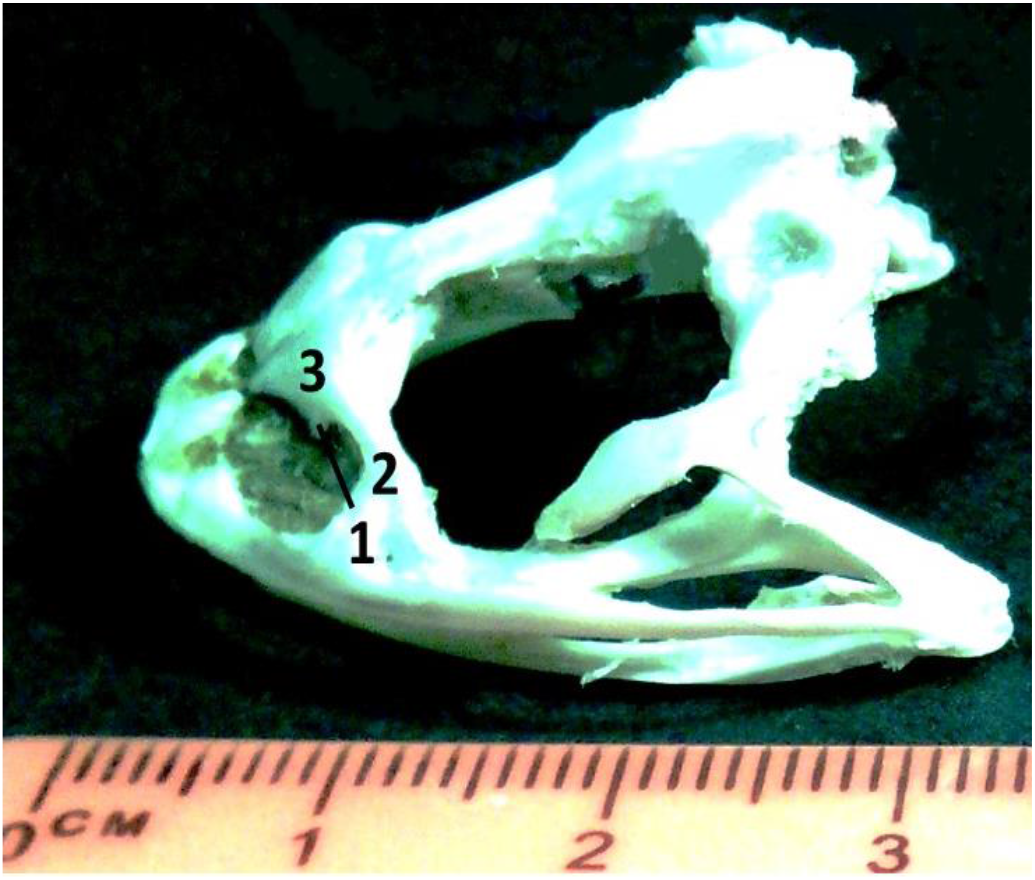
« Spike-like» protrusion on left *nasale*. Adult *P. ridibundus*. 1 – «spike-like» protrusion; 2 – *ramus lateralis*; 3 – left *nasale*.

### Upper jaw *(maxilla)*

When we studied frog *maxilla* of the genus *Pelophylax*, we found that the maxillary *(maxillare)* and quadratojugal *(quadratojugale)* bones included in its composition, at the contact region in all studied individuals overlap on each other approximately at 3 mm. This had discovered by separating these bones from each other using thin dissecting needle. The anterior end of the *quadratojugale* appear pointed and chamfered in place of contact with *maxillare*. For example, a similar junction of *maxillare* and *quadratojugale* was found in *Phyllomedussa sauvagii* (Anura, Hylidae), but in these frogs the extent of contact region of these bones varies from 1/3 to 1/2 of the length of *quadratojugale* (Ruiz-Monachesi M. R. et al., 2016). In late Cretaceous anuran *Hungarobatrachus szukacsi* Szentesi & Venczel, 2010 was found that dorsolingual surface of posterior part of *maxillare* had a shallow, triangular facet for contact with the *quadratojugale* (Venczel M. et al., 2021). In *Gastrotheca gemma* sp. nov. (Anura, Gastrotheca) from northeastern Peru the *quadratojugale* is broadly overlapped laterally by the *maxillare* (Venegas P.J. et al., 2021).

During preparation *quadratojugale* and *maxillare* of representatives of the genus *Pelophylax*, we revealed that thin oral end of *quadratojugale* passes into short connective tissue ligament by means of which *quadratojugale* is attached to *maxillare* from the inside of corresponding branch of *maxilla* (Fig. 4). *Quadratojugale* appears in tailless amphibians (Ranidae) at the end (G.R. de Beer, 1937) or immediately after metamorphosis (Roček Z., 2003). G.R. de Beer (G.R. de Beer., 1937) considered that *quadratojugale* ossifies on the outer surface of the part of square cartilage (*pars quadrata palatoquadrati*), involved in the articulation with the lower jaw, and grows forward until contact with *maxillare*. Some researchers suggest that the *quadratojugale* of adult specimens (Anura, Ranidae) appears in result of the invasion of *pars quadrata palatoquadrati* in *quadratojugale*, and this combination known as *quadratojugale* (Roček Z., 2003). In 1892 year Haupp suggested that in the representatives of Ranidae *quadratojugale* arises in result of ossification of connective tissue ligament *(ligamentum cornu-quadratum laterale)* between *maxillare* and *pars quadrata palatoquadrati,* lying laterally from the masticatory muscles (Roček Z., 2003). According to Sedra S.N. (1950), *quadratojugale* begins to develop from osteoblasts invading the rear part of the *ligamentum cornu-quadratum laterale*, not far from its attachment to the *pars quadrata palatoquadrati*. In accordance with other data *quadratojugale* develops by endesmal ossification and merges with *pars quadrata palatoquadrati* (Stephenson E.M., 1951). In representatives of Ranidae, *quadratojugale* continuously connected with partial perichondral ossification on the lateral surface of *pars quadrata palatoquadrati*, which is probably part of a square bone (*quadratum*) (G.R. de Beer, 1937). Some researchers consider that *quadratojugale* has an enlarged head at the caudal end, resulting from the ossification of *pars quadrata palatoquadrati* (Terentyev P.V., 1950). There is also records that *quadratojugale* is formed due to the ossification on caudal end of the palatine-square cartilage *(palatoquadratum)* (Gurtovoi N.N. et al., 1978). We consider that the *quadratojugale* of the representatives of *P. ridibundus* and *P. lessonae* is an integumentary bone and it developed by endesmal ossification. During of *maxilla* formation, due to the strengthening of the zygomatic arch, *quadratojugale* merges with the lateral part of the *pars quadrata palatoquadrati*. The lateral part of the *pars quadrata palatoquadrati* ossifies too (Fig. 3).

**Fig. 3.**
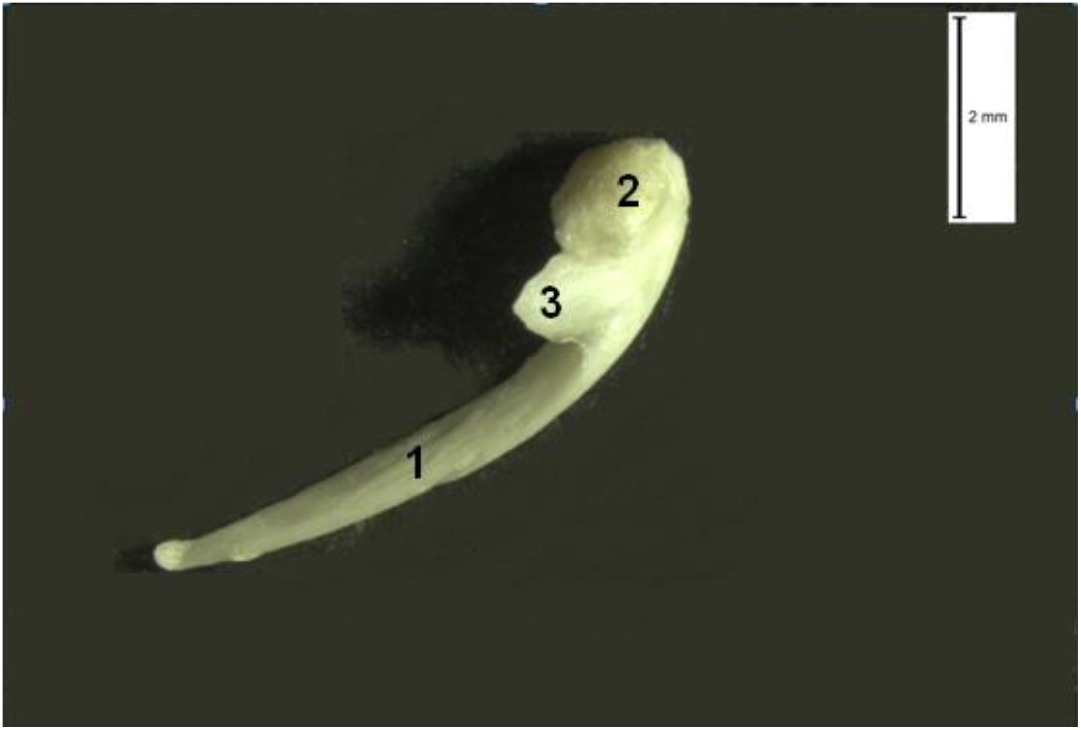
*Quadratojugale* merged with lateral ossified part of *pars quadrata palatoquadrati.* “Scaly” process of *quadratojugale*. Adult *P. lessonae*. 1 - *quadratojugale;* 2 - lateral ossified part of the *pars quadrata palatoquadrati;* 3 - “scaly” process of *quadratojugale*.

**Fig. 4.**
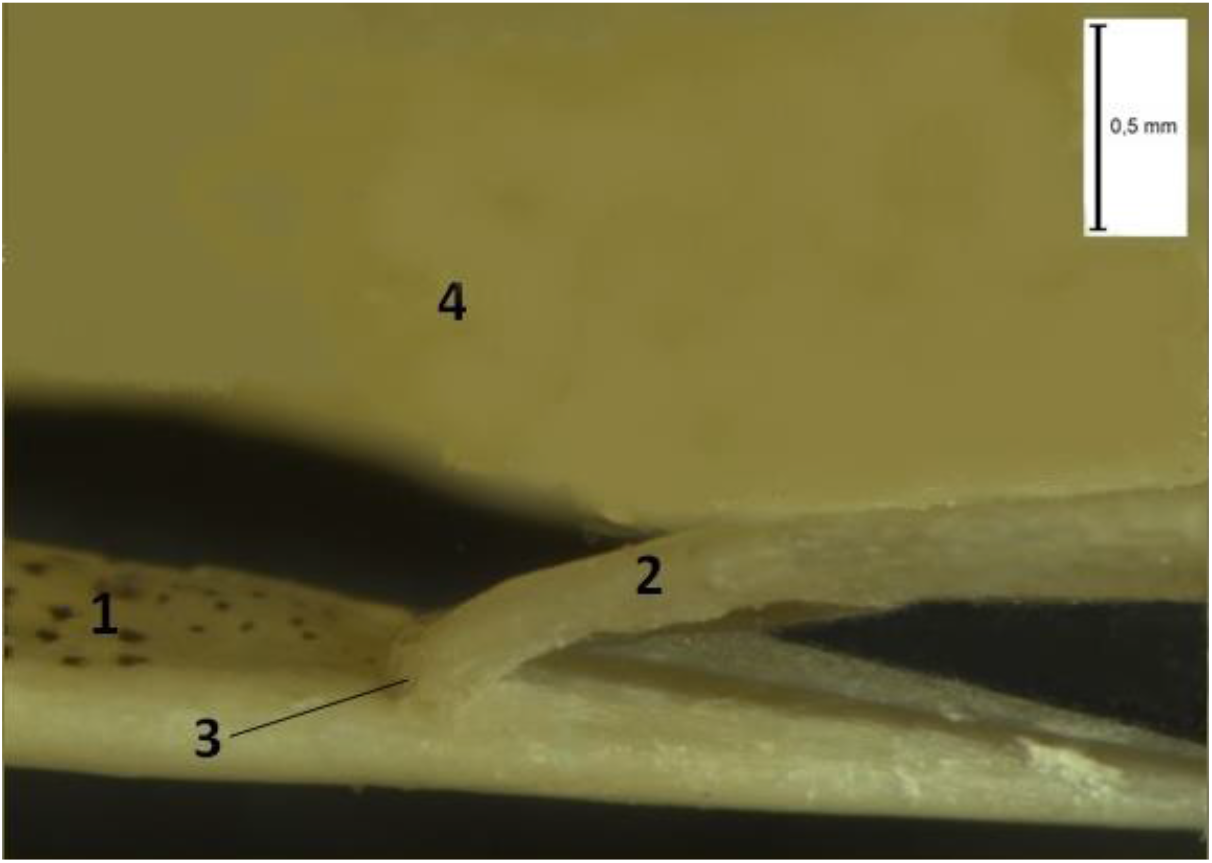
Connective tissue ligament between *maxillare* and oral end of *quadratojugale*. Adult *P. ridibundus*. 1 – *maxillare*; 2 – *quadratojugale*; 3 - connective tissue ligament between *maxillare* and oral end of *quadratojugale*; 4 – *pterygoideum*.

It is important fact, that in *maxilla* of caudate amphibians (Urodela) the absence of *quadratojugale* had found, and *maxillare* connects to *quadratum* with a long connective tissue ligament (Gurtovoi N.N. et al., 1978; Salomatina N.I., 1982; Dzerzhinsky F.Ya., 2005). The above data give us reason to consider that the preserved connective tissue ligament between *quadratojugale* and *maxillare*, revealed in frogs of the genus *Pelophylax* (Fig. 4), is homologous to the connective tissue ligament between *maxillare* and *quadratum* in representatives of Urodela. This suggests the existence of similar stages of evolutionary transformations of the skull in modern representatives of Batrachians. Homology of connective tissue ligament between the *maxillare* and the components of zygomatic arch as well as jaw joint, located more caudally in Anura and Urodela, has confirmed by following additional arguments: data for the presence of ossified part of *pars quadrata palatoquadratum* in Ranidae – part of the *quadratum* (G.R. de Beer, 1937); the fact had established by a number of researchers, that representatives of some taxonomic groups of Anura do not develop *quadratojugale*, as well as representatives of Urodela. The taxonomic groups of Anura that do not develop *quadratojugale* include *Ascaphus* and *Leiopelma* (Slabbert GK, 1945), *Pipa pipa* (Trueb L., 1989), *Xenopus* (Paterson NF, 1939), *Scaphiopus* and *Spea* (Roček Z., 1981). The absence of this bone in a number of representatives of Anura indicates that *quadratojugale* is not permanent structural element in the skull of extant species of tailless amphibians.

Many authors describe *quadratojugale* of the representatives of Ranidae, and in particularly of the genus *Pelophylax*, as rod-like thin bone (Terentyev P.V., 1950; Gamble F.W., 1999; Roček Z., 2003; Kotpal R.L., 2009). We detected, that largest part of *quadratojugale* of these frogs represents thin, slightly curved bone rod really. Closer to caudal end the *quadratojugale* expands slightly and near the confluence with ossified part of the *pars quadrata palatoquadrati* it has a wide flat process. That process tightly adjoins to caudal end of main part of the *squamosum* on medial side. We called it “scaly” process (Fig. 3). The “scaly” process didn’t describe in other representatives of Anura. *Temnospondyli* or more specifically amphibamid dissorophoids is ancestral forms of Anura and Urodela (Carroll R., 1992; Schoch R.R., 2014; Schoch R.R., 2019; Atkins J.B. et al., 2019). Apparently, *quadratojugale* and *squamosum* overlapped with each other in amphibamid dissorophoids still, and this structural feature had retained in the skull of studied frogs of the genus *Pelophylax* and serves to consolidate of bony structures of the skull and strengthen of jaw joint.

### Jaw joint

During of life the frogs of the genus *Pelophylax* retain the cartilaginous structures of the primary upper and lower jaws – palatin-square cartilage *(palatoquadratum)* and Meckel cartilage *(cartilago Meckeli)*, respectively (Gurtovoi N.N. et al., 1978; Kartashev N.N., Sokolov V.E., Shilov I.A., 2004; Kotpal R.L., 2009). These cartilaginous structures form the jaw joint (articulation). An important structural element of the jaw articulation is also the caudal part of *palatoquadratum* or *pars quadrata palatoquadrati*, which ossified in the lateral region (G.R. de Beer, 1937). According to some authors (Schmalhausen I.I., 1935; Romer A., Parsons T., 1992; Kovtun et al., 2005) in recent amphibians the caudal part of *palatoquadratum* completely ossifies and takes over the function of suspension of lower jaw. According to G.R. de Beer (1937), in the representatives of the genus *Pelophylax* caudal part of *palatoquadratum* is only partially ossified. For our data, this ossification merges with the cartilaginous fragment of caudal part of *palatoquadratum*, i.e. with a cartilaginous part of *pars quadrata palatoquadrati* (Fig. 5).

**Fig. 5.**
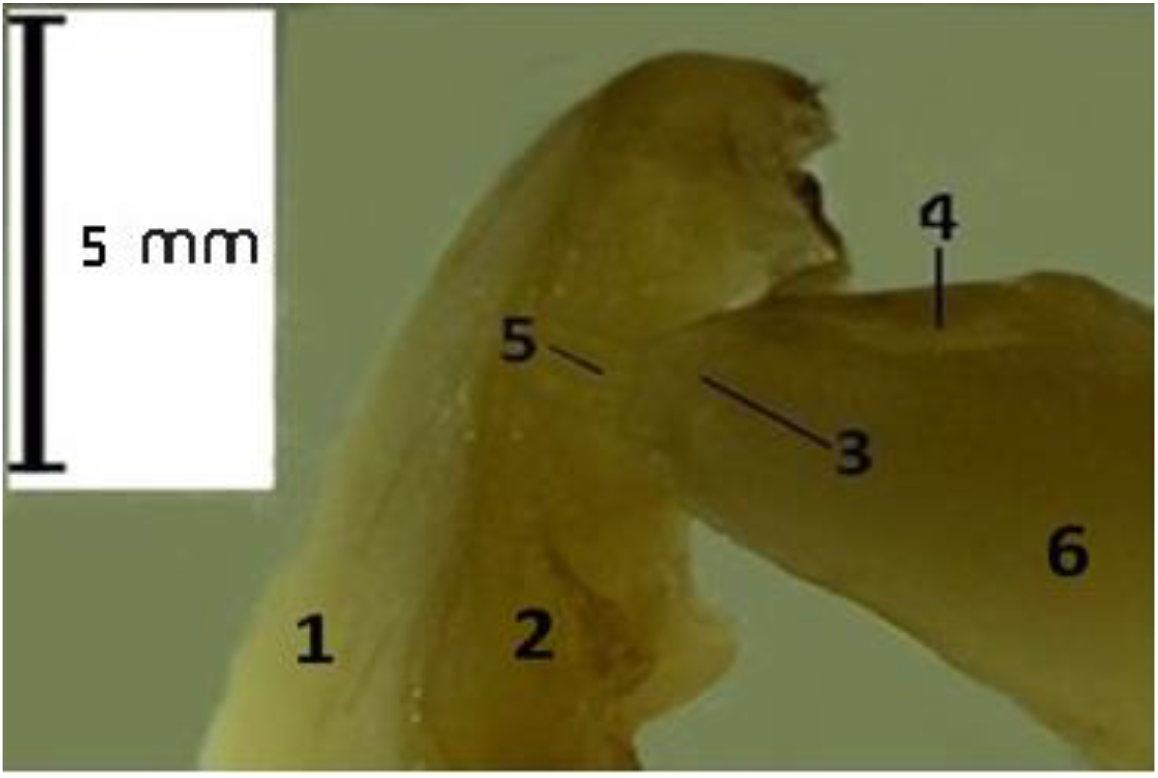
Jaw articulation of adult *P. ridibundus*. The joining point of *pars quadrata palatoquadrati* slightly offset from the caudal end of the cartilago Meckeli. 1 *– angulospleniale*; 2 – *cartilago Meckeli*; 3 – lateral ossified part of *pars quadrata palatoquadrati*; 4 – cartilaginous part of *pars quadrata palatoquadrati*; 5 – ligaments between *cartilago Meckeli* and the ossified as well as cartilaginous parts of *pars quadrata palatoquadrati*; 6 – *quadratojugale*.

We recognized that most part of the wide connective tissue ligament connected with ossified fragment of *pars quadrata palatoquadrati*, the smallest part of this ligament is attached to remaining cartilaginous fragment of *pars quadrata palatoquadrati*. The lower end of this ligament attached to the *cartilago Meckeli*. This ligament provides the necessary strength of jaw joint. Besides, under closing the mouth, this wide and strong ligament is in extended state and helps to close the jaws tightly. This is necessary for functioning of pressure mechanism of frog respiration (Bannikov A.G. et al., 1985). The second, finer connective tissue ligament, connecting the components of the jaw joint, - the ossified fragment of *pars quadrata palatoquadrati* and *cartilago Meckeli*, is located on the side of the oral cavity. It is in a stretched condition, when the frog mouth is open and serves for strengthen the jaw joint also. According to some authors in jaw joint of frogs of the genus *Pelophylax pars quadrata palatoquadrati* contacts with articular surface on caudal end of the bone of most rear part of lower jaw, *angulospleniale* (Gurtovoi N.N. et al., 1978; Kotpal R.L., 2009). Based on our observations, *angulospleniale* does not participate in the formation of jaw joint of green frogs; on the contrary, jaw joint includes *cartilago Meckeli*.

When studying the structure of jaw joint in frogs of the genus *Pelophylax*, we did not find any articular surfaces, flat sites or recess on the *cartilago Meckeli*, as well as on cartilaginous and ossified parts of the *pars quadrata palatoquadrati* involved in the formation of jaw joint (Fig. 5). By preparing green frog skulls, it can be seen that the components of the jaw joint are connected to each other only with help of connective tissue ligaments and that the jaw joint in this case does not even have a typical structure for vertebral joints. Therefore, we propose to call jaw joint in frogs of the genus *Pelophylax* by jaw articulation, like Webster D. and Webster M. (1974).

As mentioned above, the frog jaw articulation of the genus *Pelophylax* form by cartilaginous *palatoquadratum* and *cartilago Meckeli*. These cartilaginous structures are also part of jaw joint of the sharks (subclass of Chondricthyes), ancient representatives of the lower vertebrates that appeared in the second half of Devonian and survived to our time. In labyryntodons (Embolomeri, Rhachitomi, Stereospondyli), caudal areas of *palatoquadratum* and *cartilago Meckeli* were completely ossified and the jaw joint (articulation) was formed by replacement bones - *quadratum* and articular bone *(articulare)*. Nonetheless, a recess was found on upper surface of the *articulare* at Embolomeri, into which during the formation of jaw joint, the convex part of the quadratum entered (Bystrov A.P., 1957). In studied modern green frogs, *articulare* does not develop and caudal site of *palatoquadratum*, i.e. *pars quadrata palatoquadrati* is not ossifies completely. Apparently, the predominance of cartilage structures in jaw articulation of frogs of the genus *Pelophylax* is a secondary state and caused by reduction in bone structures. This decrease of the number of bone structures could occur due to the need to increase of jaws mobility during breathing and capture prey.

From the side of oral cavity *pars quadrata palatoquadrati* covered by articular part of the pterygoid bone *(pars articularis)* and from outer lateral side and partially from dorsal side *pars quadrata palatoquadrati* covered by vertical part *(corpus)* of *squamosum*. In addition, we found that the site of attachment of *pars quadrata palatoquadrati* through the ligament slightly offset from the caudal end of *cartilago Meckeli* in oral direction (Fig. 5). The caudal areas of both branches of *cartilago Meckeli* spread apart slightly to the sides, distally to *pars quadrata palatoquadrati*. Thus, the amplitude of mouth opening increases and the possibility of damage during movement of the parts of jaw joints minimize.

### Morphofunctional features of the pterygoid *(os pterygoideum)*

This integumentary bone is clearly visible ventrally and dorsally (Terentyev P.V., 1950), but it is also clearly visible from lateral side. It plays an important integrating role in the skull of frogs of the genus *Pelophylax*, linking the cartilaginous brain skull, visceral and integumentary components of the skull into a single structural complex. *Pterygoideum* consists of three bone branches: the pterygoid process *(pr. pterygoideus)* passes forward over the *maxillare*, tightly adjoining and strengthening it. The articular part *(pars articularis)* directs back and out, strengthening and protecting the *pars quadrata palatoquadrati* and jaw joint. The small basic process *(pr. basalis)* directs to parasphenoid *(parasphenoideum)* and cartilaginous auditory capsule, joins with small cartilage to auditory capsule and resembles a tube, bisected longitudinally in half (Fig. 6).

**Fig. 6.**
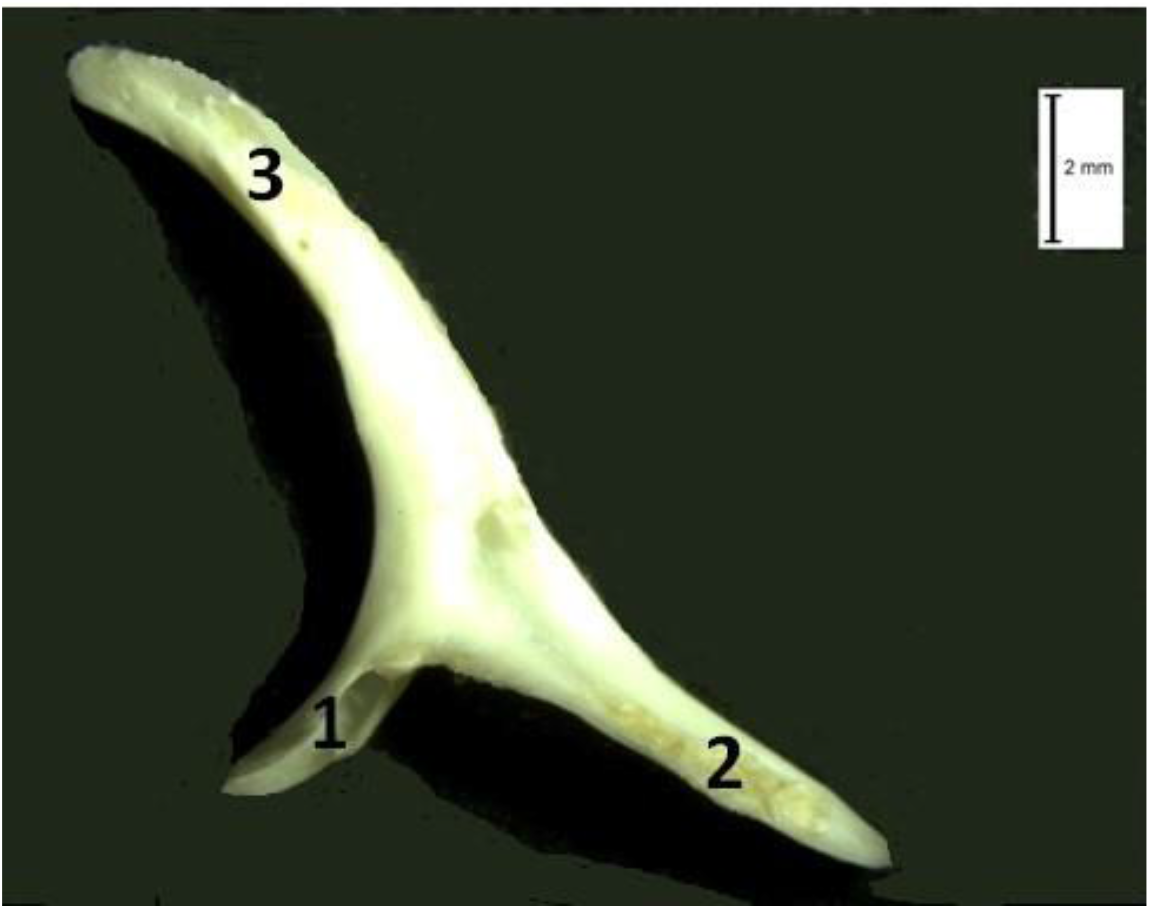
Right *pterygoideum* with *pr. basalis* in the form of a tube cut longitudinally in half. Adult *P. ridibundus.* 1 - *pr. basalis*; 2 - *pars articularis*; 3 - *pr. pterygoideus*.

A number of authors describe the location of *pr. pterygoideus* differently. There is evidence in the literature that *pr. pterygoideus* is directed forward to the *maxillare* (Terentyev P.V., 1950; Gurtovoi N.N. et al., 1978) or only contacts with back part of the *maxillare* (Roček Z., 2003). Our data do not coincide with above observations. We has found that *pr. pterygoideus* runs along the entire length of *maxillare*, over it from the side of oral cavity, tightly ajoining to *maxillare*, but not merging with it. We agree with those authors who, like us, have noted that *pr. pterygoideus* can reach the palatine bones *(palatinum)* (Gamble F.W. 1999; Kotpal R.L., 2009). We also revealed that *pr. pterygoideus* represents bone plate as if folded in half. There is the bend on the side of oral cavity, and on outside there is an unclosed cavity between two bent parts of the bone plate. Fold *pr. pterygoideus* (Fig. 7) localize precisely in area of the palate where the bone *transversum* was located at the amphibamid dissorofoids. Therefore, we suggest that the presence of this fold is the result of fusion of *pterygoideum* and *transversum* due to the formation of extensive interpterigoid windows characteristic for modern frogs of the genus *Pelophylax*. *Transversum* - the bone that was part of the palate of *Temnospondyli*. Interpterigoid windows were already present in ancestors of Anura, amphibamid dissorofoids, which lived in the Permian period (Bystrov A.P., 1957; Atkins J.B. et al., 2019).

**Fig. 7.**
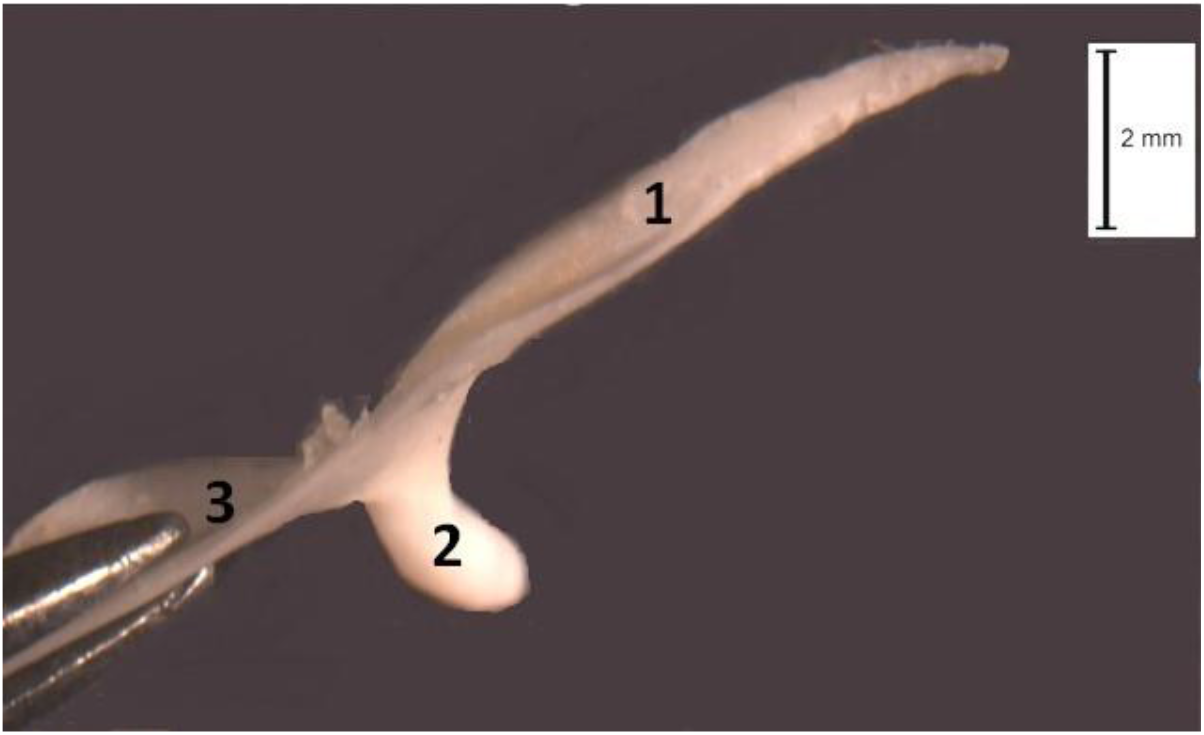
Fold*pr. pterygoideus.* Left*pterygoideum*. Adult *P. lessonae*. 1 - fold*pr. pterygoideus*; 2 - *pr. basalis*; 3 - *pars articularis*.

*Pr. basalis* forms part of posterior wall of the orbit (Kotpal R.L., 2009). We found that *pr. basalis* in green frogs, in the form of longitudinally cut tube, encompasses proximal part of small cartilage. The small distal portion of this cartilage is not encompasses by *pr. basalis* and directly adjoins to the cartilaginous auditory capsule. Based on paleontological data, we assume that this small cartilage is the remainder of the basisphenoid *(basisphenoideum)*, i.e. basipterygoid process *(pr. basipterygoideus)*. *Basisphenoideum* is the bone that lay at the base of the brain skull of the carboniferous *Temnospondyli* (Bystrov A.P., 1957). In the ancestral form of Anura, *Temnospondyli*, (Carroll R.L., 1992, 2007; Schoch R.R. 2014), *basisphenoideum* was already cartilaginous, but still maintained contact with *pterygoideum* by *pr. basipterygoideus* (Bystrov A.P., 1957). In the frogs of the genus *Pelophylax* from *basisphenoideum* only small cartilaginous *pr. basipterygoideus* retained, and function of protecting and strengthening of cartilaginous brain skull completely crossed over to parasphenoid *(parasphenoideum)*.

Some authors describe the basipterygoid joint in representatives of Anura as contacts *pr. basalis* with lateral outgrowths of *parasphenoideum* on both sides respectively (Dzerzhinsky F.Ya., 2005; Kotpal R.L., 2009). There is evidences that *pr. basalis* in representatives of Anura reaches the lower surface of auditory capsule (Roček Z., 2003). During the investigation of skull anatomy *Barbourula busuangensis* (Taylor and Noble, 1924) *pr. basipterygoideus* has been described as a process of a cartilaginous auditory capsule, which ossifies in adults (Roček Z. et al., 2016).

According to our results, the basipterygoid articulation in the frogs of genus *Pelophylax* can be described as the movable contact of small oblong cartilage, i.e. *pr. basipterygoideus*, covered *pr. basalis*, with auditory capsule. We found that there is special area on auditory capsule, which adjoins *pr. basipterygoideus*. In addition, *pr. basipterygoideus* is associated with cartilaginous auditory capsule by connective tissue ligaments. We consider that the movable cartilaginous basipterygoid articulation facilitates the movement of eyeballs in green frogs.

### Lower jaw (mandibula)

Each half of the mandibular arch of frogs of the genus *Pelophylax* consist of element of the primary lower jaw, *cartilago Meckeli*, and integumentary bones covering it - lamellar-angled bone *(angulospleniale)* and tooth bone *(dentale)*. In front of, at the place of junction of right and left half of lower jaw arch there are paired replacement chin-jaw bones *(mento-mandibulare)* (Fig. 8). We see on the outer lateral side of each half of lower jaw, that the rostral end of *dentale* overlaps on the *mento-mandibulare* and merged with it without any noticeable border between them (Fig. 9). The rest of *dentale* covers almost the entire front half of the *cartilago Meckeli*. Besides, we noticed that the groove for *cartilago Meckeli* on the outer lateral side of the *angulospleniale* has the shape of almost regular right angle, and this feature is probably due to reduction of this bone. Only at caudal end of *angulospleniale* the groove for *cartilago Meckeli* becomes more acclivous. From outer lateral side of lower jaw, *mento-mandibulare* doesn’t contact with *angulospleniale*. On the medial side of lower jaw *mento-mandibulare* and *angulospleniale* are connect to each other through connective tissue ligaments.

**Fig. 8.**
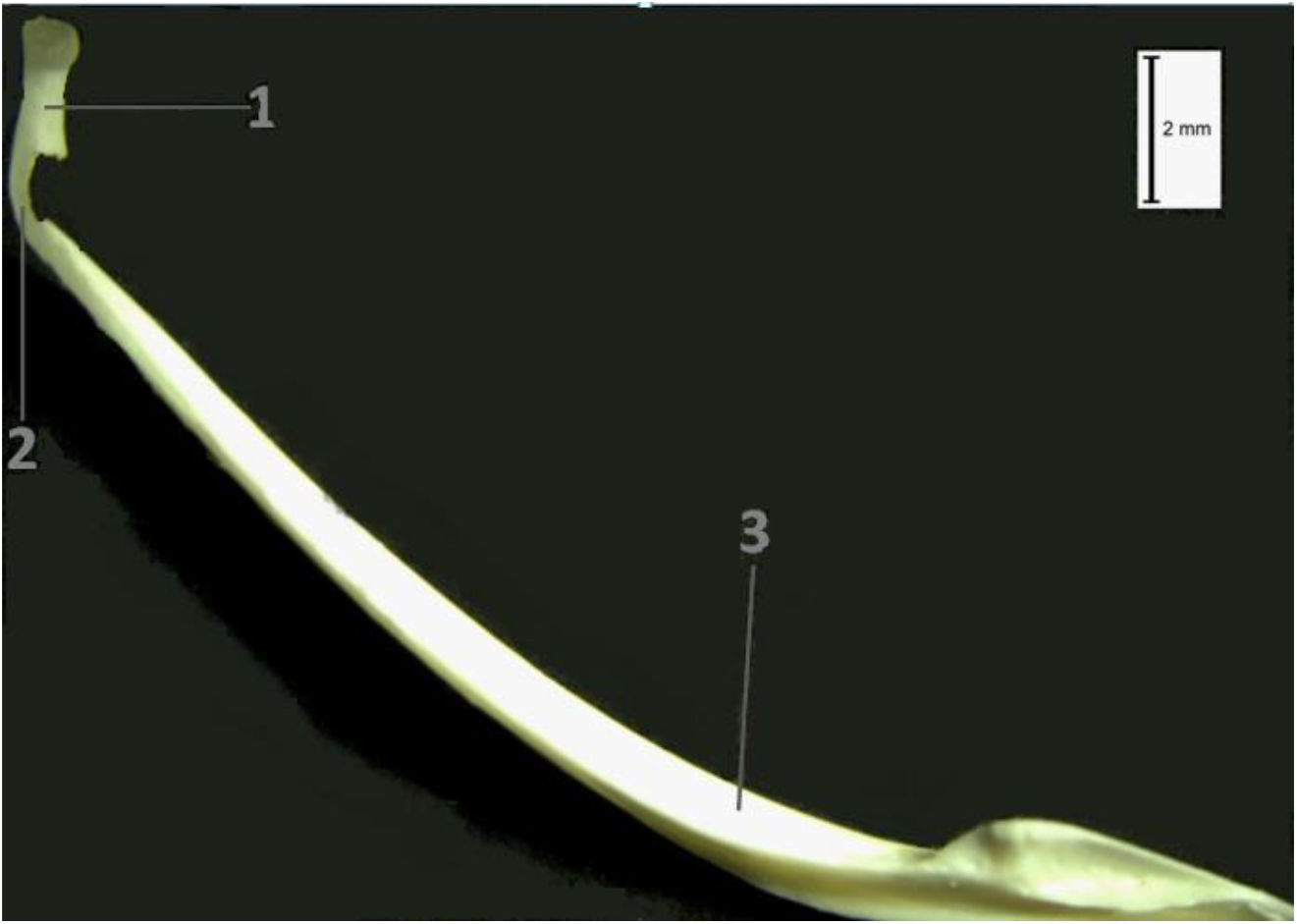
The skeleton of the left half of *mandibula*. Adult *P. ridibundus*. 1 - *mento-mandibulare*; 2 - *dentale*; 3 - *angulospleniale*.

**Fig. 9.**
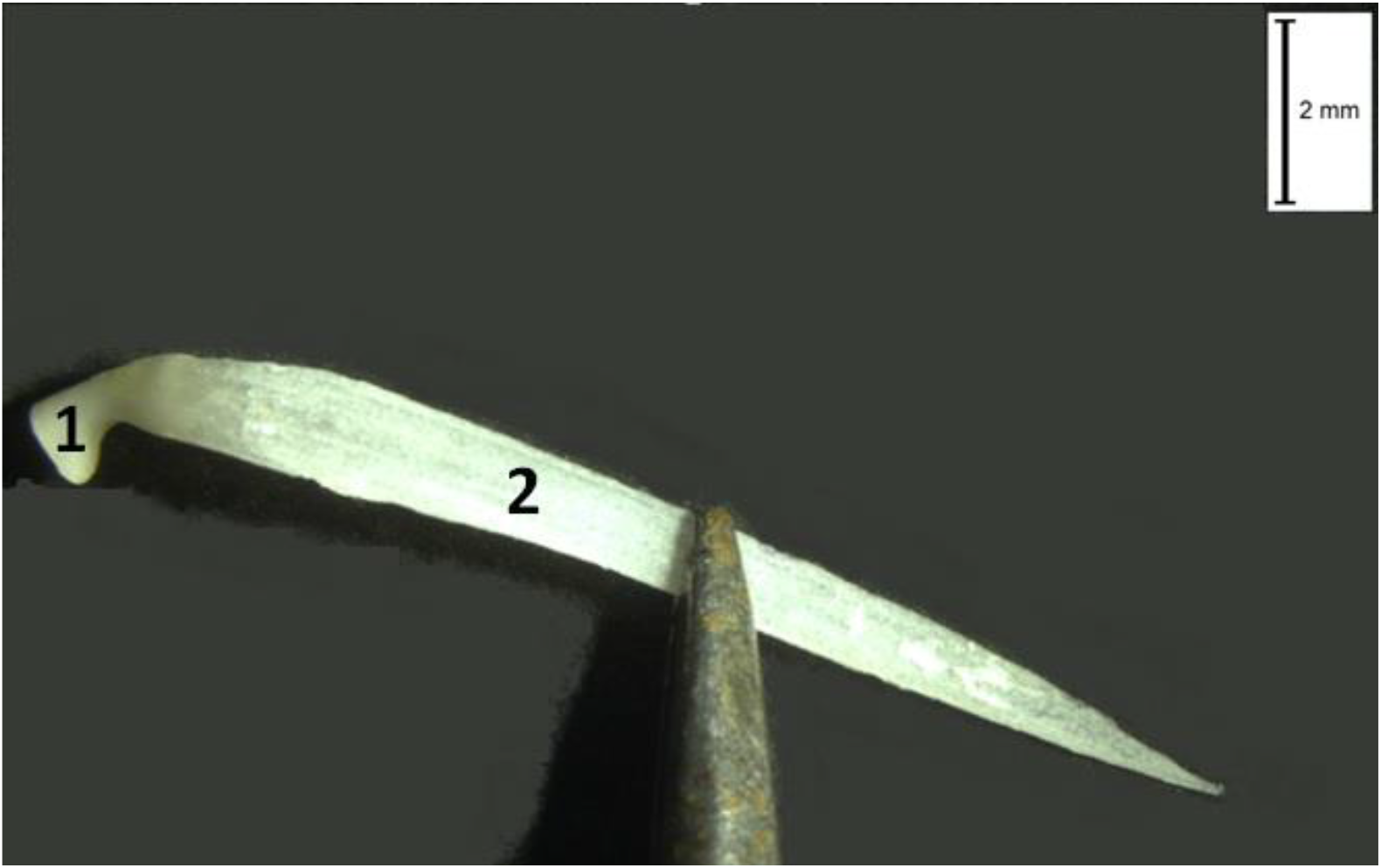
Merged *mento-mandibulare* and *dentale*. Right side of the *mandibula*. Adult *P. lessonae*. 1 – *mento-mandibulare*; 2 – *dentale*.

According to some data (Gurtovoi N.N. et al., 1978; Dzerzhinsky F.Ya. et al., 2013), there is only one unpaired *mento-mandibulare* in lower jaw of green frogs. According with data of other investigators (Yildirim E., Kaya U., 2016) and our results, representatives of the genus *Pelophylax* had paired *mento-mandibulare*. Many researchers that descript skull anatomical structure of individual representatives of the genus *Pelophylax* not call the largest integumentary bone of lower jaw a lamellar-angular bone i.e. *angulospleniale*, but call it angular bone (*angulare*) (Roček Z., 2003; Kartashev N.N. et al., 2004; Dzerzhinsky F.Ya., 2005; Konstantinov V.M., Shatalova S.P., 2005; Dzerzhinsky F.Ya. et al., 2013). Some authors called it the articular bone (*goniale*) (Terentyev P.V., 1950; Gurtovoi N.N. et al., 1978), as well as dermal articular bone (*dermaticulare* or *operculo-angulare*) (Terentyev P.V., 1950). Some researchers consider also that in lower jaw of green frogs, the bones of *angulospleniale*, *dentale* and *mento-mandibulare* are located one after another and articulate with each other (Terentyev P.V., 1950; Gurtovoi N.N. et al., 1978; Kartashev N.N. et al., 2004) that is not confirm by our data. We suggest that *angulospleniale* arose in result of the reduction and fusion of numerous integumentary bones, which were part of lower jaw of the Temnospondyli: *angulare*, *supraangulare*, *prearticulare*, *spleniale*, *postspleniale,* one or several coronoids. This is can be confirmed by the fact that *angulospleniale* is located in green frogs, as *angulare*, *prearticulare, spleniale, postspleniale* and coronoids were located in Temnospondyli (Bystrov A.P., 1957). Therefore we share the opinion of those authors, who call this bone *angulospleniale* (G.R. de Beer, 1937; Gamble F.W., 1999; Carroll R.L., 2007; Kotpal R.L., 2009; Hoyos J.M. et al., 2012; Ruiz-Monachesi M. R.et al., 2016).

*Dentale* reduced in frogs for example as in *Phyllomedusa sauvagii* (Anura, Hylidae) (Ruiz-Monachesi M. R. et al., 2016). We found that in studied representatives of the genus *Pelophylax dentale* had undergone reduction too. Reduction of *dentale* might occurred due to shorthening of the lower jaw in frogs (Paluh et al., 2021). Shorthening of the lower jaw is a skeletal trait that is known to occur in species that catch and eat small prey (Vidal-Garcia M, Scott Keogh J., 2017; Paluh et al., 2020), in particularly green frogs of the genus *Pelophylax. Dentale* of green frogs are thin bony plates, shortened, tapering to their caudal end and barely reaching half the length of *cartilago Meckeli*. The necessary and main tool for prehension of prey in frogs of the genus *Pelophylax* is secondary upper jaw (Dzerzhinsky F.Ya., 2013). Functional role of the upper jaw, as well as, probably, ensuring tight closing of the mouth during operation of pumping mechanism of respiration caused the reduction of *dentale* of green frogs. Green frogs have no teeth on the lower jaw. Perhaps reliable closing of the mouth during respiration of green frogs is also promote the shape of groove for *cartilago Meckeli* and the fact, that dorso-lateral side of posterior half of *cartilago Meckeli* not covered by any bones and it is directly under the skin.

The structure of lower jaw of the frogs of the genus *Pelophylax* has specializated features associated with its functional role in the organizm: ensuring the operation of pumping mechanism of respiration, exit from the catching prey.

## Conclusions

1. The features of frog skull anatomy of the genus *Pelophylax*, that became the subject of present study, found in all investigated individuals of species *P. ridibundus* and *P. lessonae*. Therefore, we can assume that they are typical for all representatives of these species.
2. The frog skull of *P. ridibundus* and *P. lessonae*, as well as in many other representatives of Anura, shows highly specific morphological features compared to the skulls of representatives of other groups of terrestrial vertebrates, for example, the presence of *mento-mandibulare* and *angulospleniale* in lower jaw, a large amount of cartilage in the skull.
3. The morphological features of frog skull of *P. ridibundus* and *P. lessonae* are adaptive and closely related to the functions of skull in the organism and with characteristics of the habitat.
4. The skull of frogs of *P. ridibundus* and *P. lessonae* retains ancient structural features inherent for ancestral form, Permian Temnospondyli.
5. The anatomical structure of skull of representatives of *P. ridibundus* and *P. lessonae* is interesting for further studies.

## Acknowledgments

The author thanks the staff of the Department of Evolutionary Morphology of Schmalhausen Institute of Zoology for making valuable suggestions on the manuscript. The author would also like to thank the staff of the Department of Evolutionary Genetic and Fundamentals of Sistematic of the same institute for providing access to the materials of herpetological collection of the department.

